# Interplay between harvesting, planting density and ripening time affects coffee leaf rust dispersal and infection

**DOI:** 10.1101/2023.10.31.564961

**Authors:** Emilio Mora Van Cauwelaert, Kevin Li, Zachary Hajian-Forooshani, John Vandermeer, Mariana Benítez

## Abstract

The relationship between the spatial structure of hosts and the individual movement of vectors is central to developing a comprehensive representation of pathogen dynamics. Here we focus on the dynamics of coffee leaf rust (*CLR; Hemileia vastatrix*), which is one of the principal coffee diseases around the globe. This fungus disperses between plants or regions through direct contact, water splash, or local turbulent wind conditions. Some studies have proposed that coffee harvesters can also bear CLR during harvesting. However, it is not clear how relevant these movements are for CLR dispersal and plot-level infection in plots with different planting densities. Besides, there are bidirectional interactions between the plantation characteristics (e.g. planting density and ripening synchronicity) and the movement of harvesters that might modify their dispersal potential. Overall, there is a lack of mechanistic models to represent these interactions. Here we present a computational model to explore the role of coffee ripening synchronicity in reproducing the different movement trajectories observed in the field. We then evaluate how these harvesting trajectories modify rust dispersal and change CLR infection in plots with increasing planting densities and a constant contact-mediated rust dispersal mechanism. For both synchronization scenarios, the distribution of step lengths is positively skewed. Interplant asynchronous ripening and scarcity of trees with berries generate trajectories with medium to long steps contrary to scenarios with synchronous ripening and without scarcity. The harvest dispersal significantly increases coffee plot infection (up to 15%) compared to scenarios where only local mechanisms for CLR dispersal are present. This effect is maximal for trajectories with medium to long steps from the asynchronous scenario and in plantations with medium planting densities. In particular, medium-sized steps are more likely to create new foci and networks of contact-mediated infected plants. The interaction between the proportion and the size of infected networks results in a nonlinear behavior of rust increase due to harvest. Our results aim to spur discussion on practices that could reduce the impact of harvesting and specific trajectories, in scenarios where one can benefit from asynchronous maturation of berries and shaded plantations. Some examples of practices include reducing the medium to long-distance movements during the same day, for instance, by avoiding harvesting at the end of the harvest season when trees ripening asynchronicity and CLR levels are higher.

## 1. INTRODUCTION

The life cycle of pathogens in plants can be divided broadly into two stages: the infection stage where the organism invades the host and reproduces, and the dispersal stage where new replicates of the pathogen move from one host to the other (Avelino et al., 2004; Pangga et al., 2011). The rate and extent at which both stages are completed largely determine the magnitude of the epidemic and the crop loss (Avelino et al., 2004). Contrary to epidemics in non-sessile organisms, the host’s density and spatial arrangement play a central role in plants, as they can modify the range and dynamics of the pathogen’s dispersal (Keeling, 1999; Vandermeer et al., 2018; Papaïx et al.; 2015). In agriculture, this spatial structure depends on management factors (Beasley et al., 2022), ecological dynamics (Li et al., 2016), and also on the topography of the field, among others (Soto-Pinto et al., 2000). Additionally, the interaction of the arrangement of the hosts and the disease depends on the characteristic transmission’s distance of dispersal (Park et al., 2001; Hajian-Forooshani and Vandermeer, 2021). Few studies compare the effect of different mechanisms of pathogen transmission across spatially explicit landscapes (White et al., 2018). Studying these spatial relationships is crucial for an ecological approach to pest management that does not rely on synthetic pesticides, or solely on resistant plant varieties that focus on the infection stage (Avelino et al; 2004).

Coffee leaf rust (CLR), caused by *Hemileia vastatrix* Berk. & Br is one of the principal coffee diseases and has caused devastating loss of production in different coffee sites around the globe (McCook and Vandermeer, 2015; Talhinhas et al., 2017). Its mechanisms for dispersal have been amply studied (Becker and Kranz, 1977; Kushalappa and Eskes, 1989; Li et al., 2023; Vandermeer et al., 2018) and thus provide a great model to explore the disease’s spatially explicit dynamics and its relations with the host’s spatial arrangement and density (Beasley et al., 2022; Gagliardi et al., 2020; Hajian-Forooshani and Vandermeer, 2021; Mora Van Cauwelaert et al., 2023a). CLR forms urediniospores that can travel to neighboring coffee plants through direct contact, water splash, or local turbulent wind conditions (Kushalappa and Eskes, 1989). The urediniospores may also ascend to the atmosphere and travel to other plantations (Becker and Kranz, 1977; Vandermeer and Rohani, 2014). For each of these ways of dispersal, there are agroecological management proposals to reduce the spread, like reducing the density or clustering of the plantations (Beasley et al., 2022; Ehrenbergerová et al., 2018), avoiding regular or extremely dense patterns that can make the epidemic thrive through contact or splash-mediated dispersal (Hajian-Forooshani and Vandermeer, 2021; Mora Van Cauwelaert et al., 2023a; Li et al., 2023), but also planting different trees within the plots and in the borders to reduce local and regional wind dispersal (Avelino et al., 2022; Boudrot et al., 2016; Gagliardi et al., 2020)

It has been observed that workers can also disperse CLR within the plot and between regions (Becker and Kranz, 1977; Ramírez-Camejo et al., 2022; Schieber and Zentmyer, 1984). At the plot scale, this relates to the fact that some foci of infection occur close to harvesting paths and human settlements (Waller, 1982, 1972) and that the build-up of the epidemic correlates with the harvesting season (Avelino et al., 1991; Mora Van Cauwelaert et al., 2023a). However, it is unclear how relevant the dispersal by harvesters is for the whole epidemic in contrast to other modes of dispersal, and what agroecological measures could be implemented to reduce its impact.

Besides, the dynamics of harvesting in coffee plantations depend on the scale, on specific ecological and management characteristics, but also on land tenure regimes and goals of production; or on what has been called “the syndrome of production” (Andow and Hidaka, 1989; Perfecto et al., 2019). In large-scale and landlord-owned plantations, the trajectory of paid workers during harvest relies, among other factors, on the ripening synchronization of the berries between the trees in the plot (Mora Van Cauwelaert et al., 2023b). Distinct trajectories, with short or long steps between plants, could play a differential role in CLR dispersal and its impact on plantations (Mora Van Cauwelaert et al., 2023b).

Here we present a mathematical model to study the workers’ spatial movement during coffee harvesting and its contribution to CLR dispersal and infection at a plot level. We first explore how the spatial trajectories drawn by the workers in large-scale plantations (Mora Van Cauwelaert et al., 2023b) could be reproduced in simulated coffee plots with different berries maturation synchronicity. We then evaluate how these trajectories modify rust dispersal and affect the final CLR infection in plots with increasing planting densities. Finally, we discuss the implications of our results in a broader context of different syndromes of coffee production.

## 2. METHODS

The objective of implementing a computational modeling approach was twofold: i) test if the temporal differences in ripening between the trees are sufficient to reproduce the reported trajectories of the workers during harvest (as previously speculated by Mora Van Cauwelaert et al., 2023b), and ii) explore the effect of these trajectories for dispersal and average rust infection in coffee plots. We first simulated coffee plots with different planting densities, interplant ripening scenarios and a closest-neighbor harvesting rule to study the role of ripening on harvesting trajectories. Then, we defined a harvesting-based mechanism of rust dispersal to evaluate the impact of harvest in the average coffee plot rust infection alongside a contact-mediated rust dispersal mechanism. In the following lines, we describe the specificities of the model (Fig. 1), the simulated scenarios, and the analyses made for each objective.

**Fig. 1.**
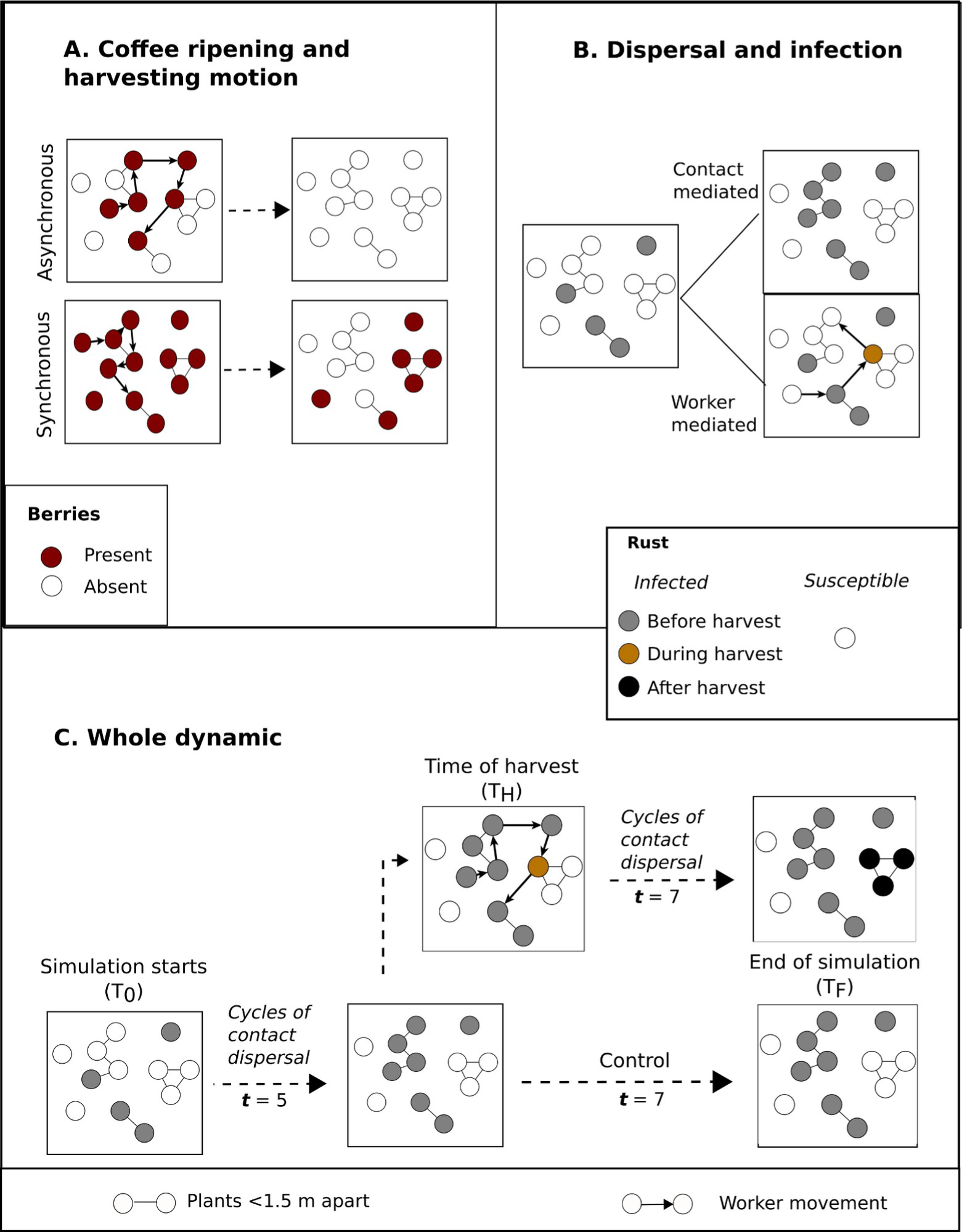
Model structure and dynamics. ***A)** Coffee ripening and harvesting motion.* We illustrate the two coffee ripening scenarios explained in the text. In the asynchronous scenario half of the plants in the plot have ripe berries (red circles) at the date of harvest, and in the synchronous scenario all the plants have ripe berries. The presence of berries determines the trajectory of the worker (black arrows) during harvest. ***B. Dispersal and infection***. Plants can be susceptible (white circles) or can be infected before the harvest (gray circles), during the harvest (yellow circles) or after the harvest (black circles). Plants can be infected through contact-mediated dispersal when they are 1.5 m apart or less from an infected plant (gray circles), or through worker-mediated dispersal (yellow circle). ***C) Whole model dynamic.*** We depict the start of the simulation (T_0_), the five cycles of contact dispersal and infection until the time of harvest (T_H_) and the two possible scenarios afterwards: the control scenario where no harvesting takes place, and the harvest scenario where harvesters can create new infections (yellow circles) that might disperse to the neighbors after new seven cycles of contact-mediated dispersal and infection (black circles).

### 2.1 Model Description

We represent one hectare coffee plots as 100 x 100 unit squares with non-periodic boundaries, with N randomly placed points representing the plants (randomized arrangements are observed particularly in old plantations; Hajian-Forooshani and Vandermeer, 2021). As we fix the area, N represents the average planting density per ha. Plants within the simulated plantation are assumed to be homogeneous, except for two variables: the ripening and the infection status.

#### Coffee ripening and harvesting motion

Each plant can have coffee berries (***present***) or not (***absent***) (Fig.1A). In order to explore the role of ripening on harvesting dynamics, we defined two coffee interplant ripening scenarios. In the asynchronous scenario (**A**), 50% randomly selected plants have ripe berries at the date of harvest. In the synchronous scenario (**S**), all the plants have ripe berries at harvest (Fig. 1A). The presence of berries defines the harvesting motion. Each worker starts harvesting in a plant with berries, collects them (changing the status of berries of that plant to ***absent***), and moves to the closest plant with berries (Fig.1A). This cycle repeats until half of the plants in the plots are visited. In the asynchronous scenario, this leads to a plot without any berries left, and in the synchronous scenario half of the plants will still have berries (Fig. 1A). According to previous studies one worker collects around 30 plants per hour, or roughly 250 plants in eight hours (i.e. one harvesting day; Mora Van Cauwelaert et al., 2023b). In the following simulations, we adjust the number of workers in the plots to always represent one harvesting day with 250 harvesting steps per worker (e.g. if we have 2000 plants, we will need four workers to visit half of the plants in a day).

#### Rust dispersal and infection

Each plant has one of two states of infection: ***susceptible*** or ***infected***. In our model we used ***infected*** to refer to plants that are infected and infective (with rust spores present). When an infected plant infects susceptible neighbors at time step ***t***, these become infective at ***t+1***. Each unit represents roughly 30 days (i.e. one month), this is, the time needed for rust to complete a cycle of infection (Leguizamon et al., 1998). An infected plant can disperse and infect the neighboring plants by two mechanisms (Fig. 1B). Firstly, they disperse to all 1.5 m apart or less susceptible neighbors through ***contact-mediated dispersal***. This mimics the local dispersion of rust either by contact between two nearby plants or by splash (as proposed by Vandermeer et al., 2018). The process creates networks of infected nearby plants where each pair of plants are 1.5 m apart or less (gray circles in Fig. 1B). Secondly, when a worker harvests an infected plant and reaches a susceptible plant in the following harvesting step, it will become infected (yellow circle in Fig.1B). We call this the ***worker-mediated dispersal***. Both mechanisms are assumed to be instantaneous in relation to time step ***t***.

#### Whole infection dynamics

To analyze the effect of harvesting in rust infection, we simulate a whole season dynamic where each plot started with an initial percentage of infected plants and got through twelve cycles of contact-mediated dispersal and infection (control scenario, Fig. 1C). This represents the maximum rust spread during one year, without any harvesting and without acknowledging for leaf loss or other mechanisms or climate factors that in practice reduce the general infection (Kushalappa and Eskes, 1989). In the harvesting scenarios, we add a punctual harvesting day event a **T_H_** after five time steps (**t** = 5). These five cycles mimic the five months from the beginning of CLR buildup to the peak of harvesting (Mora Van Cauwelaert et al., 2023a). At the date of harvest (**T_H_**), almost all the plants in the same contact-mediated networks with one or more initially infected plants are infected (gray circles in Fig. 1C). During harvest, new infections emerge (yellow circles in Fig. 1C). After harvest, seven new cycles of contact-mediated dispersal and infection take place in which the newly infected plants spread the rust across their contact-mediated networks (black circles in Fig.1B).

### 2.2. Scenarios and measurements

We first characterized the trajectories and the distribution of steps simulated under two scenarios of coffee ripening (A and S) and five planting densities (500, 1000, 2000, 3000, and 5000 plants per ha; Soto-Pinto et al., 2000), adding up to 10 scenarios. We replicated each scenario 30 times to account for the random initial arrangement of plants. We compared the trajectories drawn in plantations with 2000 plants/ha with published data of plantations with equivalent densities and different levels of ripening synchronicity (Mora Van Cauwelaert et al., 2023b). In particular, we took two examples from a large-scale organic plantation with an asynchronous ripening at the date of harvest (“organic plantation”, henceforth) and two from a slightly shaded, large-scale non-organic plantation with synchronous ripening at the date of harvest (conventional plantation). Both plantations had a planting density between 2000 to 3000 plants/ha. Secondly, we used our model to explore the effect of these simulated trajectories for each coffee ripening condition and planting density, on the plot average rust infection. We ran each scenario 30 times for the whole season (**t= 12**) starting with 20% of initially infected plants and compared them to non-harvesting scenarios (five additional scenarios, one for each planting density; Fig.1C). Finally, to further explore the mechanisms behind our results, we registered the networks of nearby plants (where each pair of plants are 1.5 m apart or less) infected through the contact-mediated dispersal mechanism before harvest (gray circles in Fig. 1B) and after the harvest event (black circles in Fig. 1B). We calculated their proportion and their mean size in terms of number of nodes and included the product of both magnitudes as a proxy for the number of infected plants after the harvest event. We also recorded the newly infected plants per step lengths during harvest (yellow circles in Fig. 1B).

The model and simulations were implemented in the *Python* 3.10.6 programming language, using the modules *NumPy, SciPy, Pandas, and Seaborn*, and ran on the LANCIS facilities, at the Ecology Institute of UNAM. The data analyses and figures were done in *Rstudio* 2023.03.1+446 using *plyr, dplyr, tidyverse, ggplot2, patchwork* libraries, and *Inkscape 1.0.* All code and data required to reproduce results in the work can be accessed at https://github.com/tenayuco/toyModel_harvest.

## 3. RESULTS

### 3.1. Interplant coffee ripening synchronization and berry scarcity produce two qualitatively different harvesting trajectories previously documented in large-scale plantations

The deterministic movement of the simulated workers across randomly distributed plants generates random trajectories (Fig. 2 and Fig.S1.1). For both synchronization scenarios, the distribution of step length is positively skewed and can be approximated with continuous lognormal and exponential distributions (Fig. 2, Fig.S1.2 and Fig. S1.3). Within each planting density, the mode is equivalent (Table S1.1). In the asynchronous ripening scenario with 2000 plants/ha, 27% of the steps are smaller than 1.5 m, 95% are below 9 m (short to medium steps), and there are some unlikely long steps (30 to 100 m) that produce a heavy-tailed distribution (Fig.2, first and second column). In the synchronous scenario, 51% of the steps are smaller than 1.5 m, 95% are below 4 m (short steps), and all steps are smaller than 30 m, creating a short-tailed distribution (Fig.2, first and second column). In the asynchronous scenario, the workers explore a larger area than in the synchronous scenario (Fig. 2, second column).

**Figure 2.**
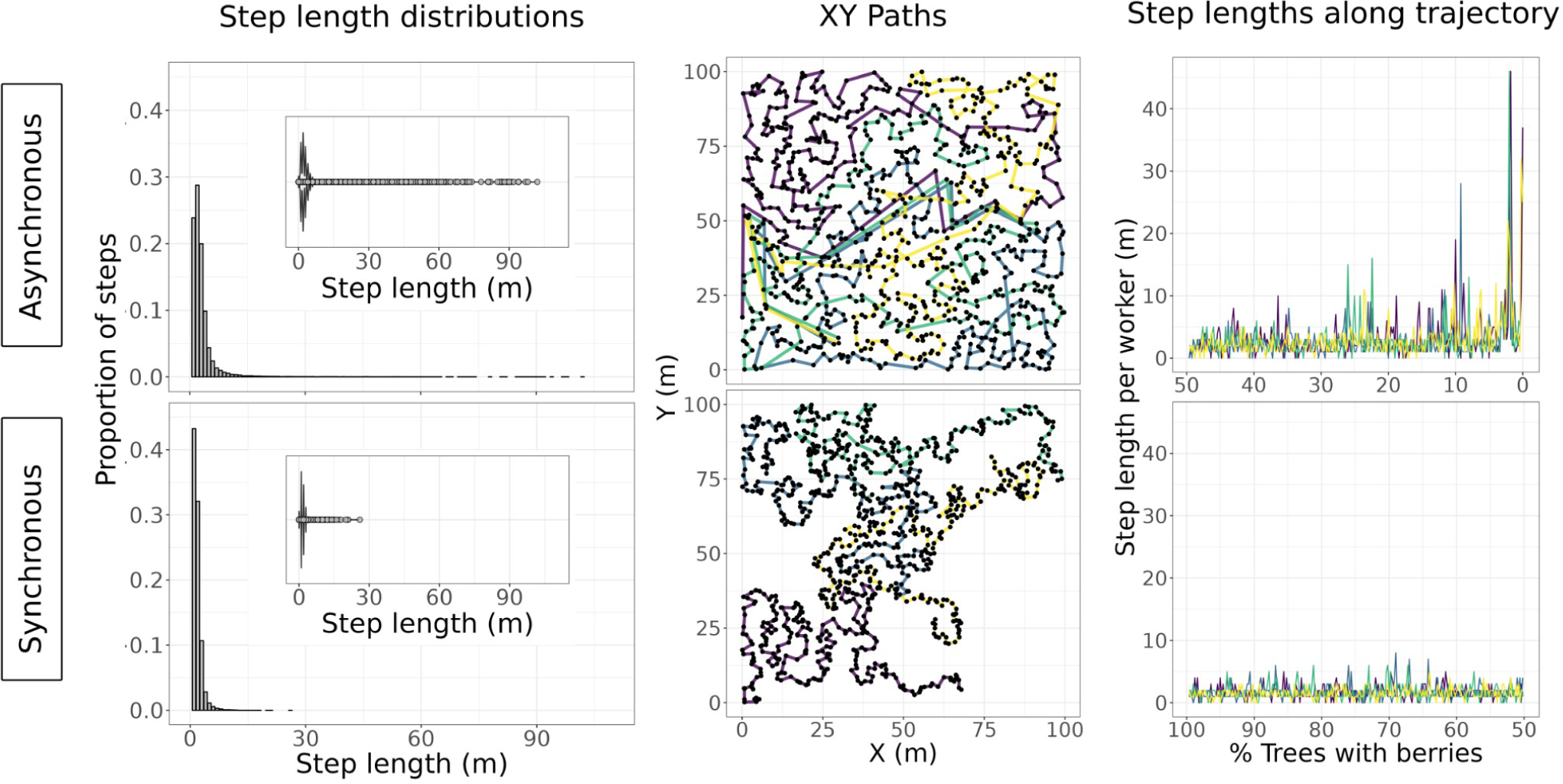
Simulated harvesting trajectories under two scenarios of coffee ripening in plots with 2000 plants/ha. We present the step length distribution (first column), the XY paths (second column), and the step lengths along the percentage of trees with berries (third column). ***The step length distribution*** summarizes the results of 30 repetitions per scenario. The boxplot was included as an aid to visualize longer steps. The modes of the skewed distributions are 1.7 m and 1.3 m for the asynchronous and synchronous scenarios, and the means are 3.2 m (± 5.2) and 1.7 m (± 1.2), respectively. For the presented data, the number of workers is 4. ***The XY path*** shows the spatially explicit movement of the workers for one of the repetitions in each coffee ripening scenario. Each color represents one worker. In ***the step length along the trajectory*** we plot the length of each step during harvesting for each worker in relation to the increasing scarcity of trees with berries due to harvest. In the asynchronous scenario, no trees with berries remain at the end of harvest, in the synchronous, half of the trees will still have berries (Fig. 1A). The trajectories and step length distributions for other planting densities are presented in Fig. S1.1, Fig. S1.3, and Fig. S1.4).

Interestingly, the presence of longer steps (>30 m) is not directly related to the initial low density of plants with berries in the asynchronous scenario. In fact they are not present in any of the equivalent and sometimes lower planting densities in the synchronous scenarios (Fig. S1.1). Long steps are concentrated near the end of the harvesting process for all planting densities, suggesting they are related to an increasing scarcity of trees with berries during the harvesting (Fig.2, third column and Fig. S1. 4).

The generated trajectories per worker at a planting density of 3000 plants/ha are qualitatively similar to in-field trajectories drawn by workers in two large-scale coffee plantations with equivalent planting density (3000 to 4000 plants/ha) but that differed in their ecological management and their ripening synchronicity at the date of harvest (Fig. 3). The modeled asynchronous trajectory is equivalent to the trajectories generated in an organic plantation with an asynchronous interplant ripening at the date of harvest (O in Fig. 3, green line). The synchronous model is qualitatively similar to the trajectories generated in a plot where ripening was more homogeneous (C in Fig. 3, blue line). Although other processes and variables may affect trajectories, this result implies that the coffee ripening differences and the scarcity of trees with berries at the end of harvesting in our model could be *sufficient* to generate the different step lengths observed in the field. In our model, when the ripening between plants is asynchronous, workers generate a higher number of medium steps to reach plants with berries, and longer steps between coffee plots when a zone is depleted (Fig. 3). Contrarily, synchronous ripening or a short harvesting process (before reaching the scarcity of trees; Fig. 2) causes workers to stay in the same coffee rows and area during the day (Fig.3).

**Fig. 3.**
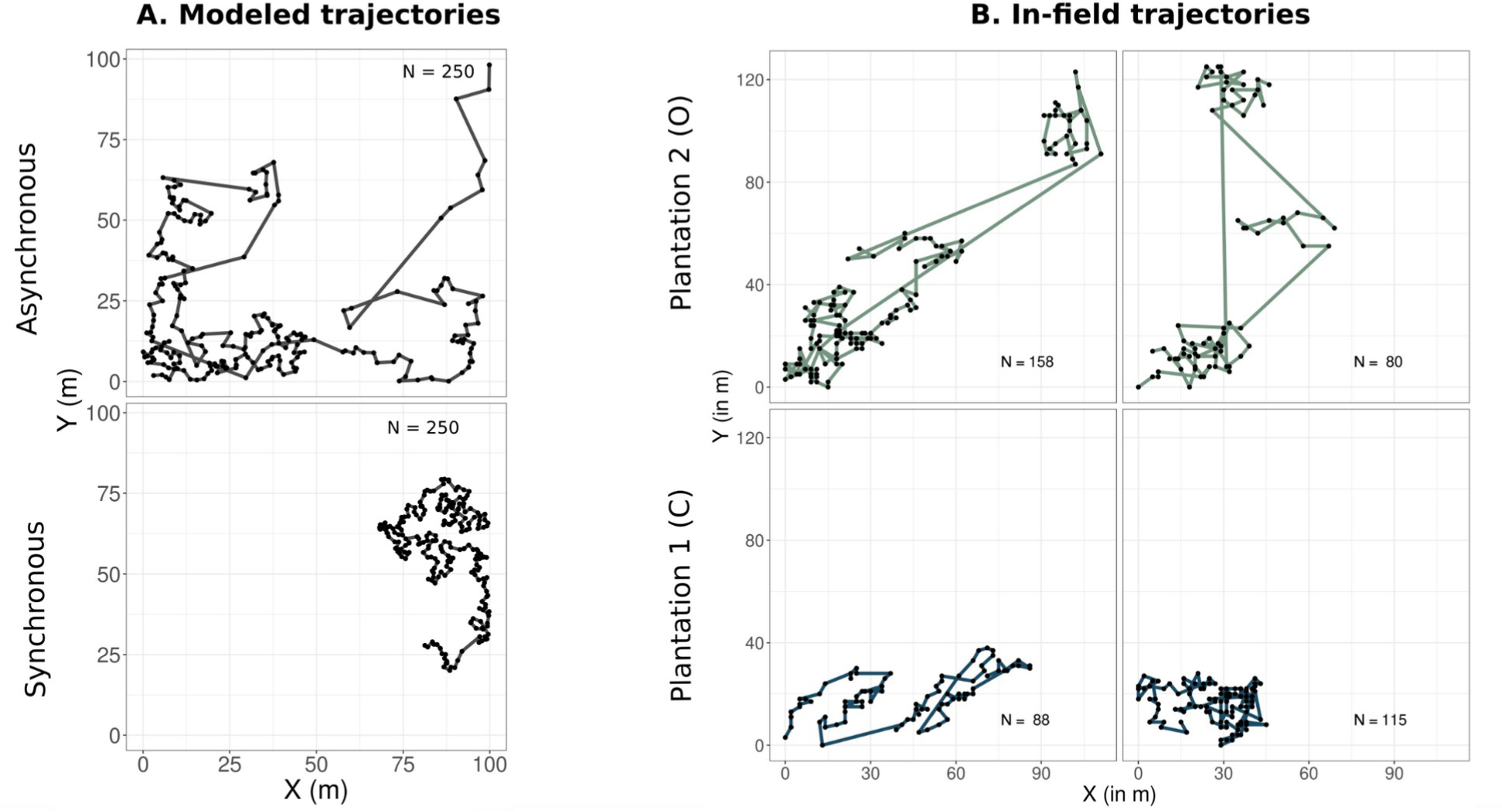
Comparison of the model with in-field trajectories. **A.** Two examples of the modeled trajectories drawn by one worker under the asynchronous and the synchronous scenarios, with N= 3000 plants/ha. **B.** Examples of in-field registered trajectories from Mora Van Cauwelaert et al. (2023b). The organic plantation (green line) is a large-scale shaded and organic plantation with an asynchronous ripening at the date of harvest. The conventional plantation (blue line) is a slightly shaded, non-organic large-scale plantation with synchronous ripening at the date of harvest. We depict the number of visited plants for each trajectory (N) (see Mora Van Cauwelaert et al., 2023b for more details). Both plantations are around 300 ha, with a planting density between 3000 to 4000 plants per ha.

### 3.2. Dispersal during harvesting increases final rust infection, especially in the asynchronous scenario and in plots with medium planting densities

In this section we analyze the effect of the different generated simulated trajectories (with medium and long steps from the asynchronous ripening scenario, and short steps from the synchronous scenario) on the average rust in the plot.

Higher planting density increases the plot-average infection in scenarios where only contact-mediated dispersal is present (control; white line in Fig. 4A). Nonetheless, for each planting density, adding the harvesting event increases the final infection compared to the control, specially in the asynchronous ripening scenario (orange and blue lines in Fig. 4A). The difference between the average rust with harvesting and without harvesting follows a non-linear curve with increasing planting density for both kinds of harvesting scenarios (orange and blue boxes in Fig. 4B). The maximum effect of harvesting in the asynchronous scenario lies at 2000 plants/ha (∼15% average rust increase; blue box in Fig. 4B), and between 1000 and 2000 plants/ha for trajectories in the synchronous scenario (∼ 8% increase; orange box in Fig. 4B). At very low densities (500 plants/ha) the difference between both scenarios is not distinct from zero. At 5000 plants/ha, the impact of harvesting drops for both scenarios as almost all the plants are already infected by contact (Fig 4.A and B). The difference between harvesting and non-harvesting scenarios remains with initial conditions from 5 to 30% of initially infected plants with maxima from 3000 to 1000 plants/ha (Fig. S1.5). At very low (1%) and very high (50%) initial infection scenarios, the effect of harvesting is more pronounced at higher densities (5000 plants/ha) and minimal densities (500 plants/ha), respectively (Fig. S1. 5)

**Figure 4.**
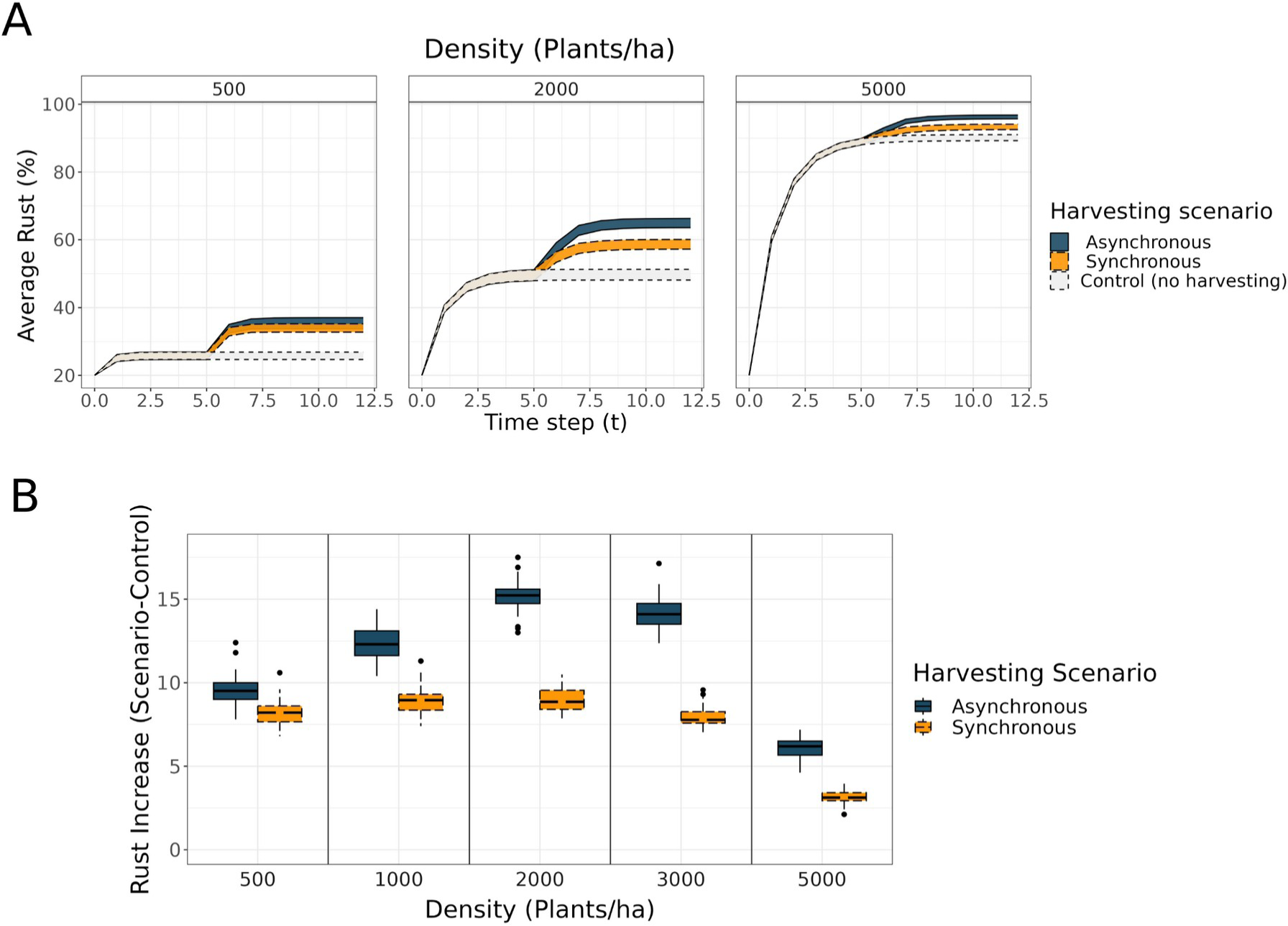
Effect of different harvesting trajectories on the average plot rust infection. ***A.*** Full time series of the average rust for three planting densities, for the control and harvesting trajectories generated under the two coffee ripening scenarios (white: control, blue: asynchronous ripening scenario, orange: synchronous ripening scenario). The width of each line represents the variability (± one standard deviation around the mean) for the 30 repetitions. The time series for 1000 and 3000 plants/ha were equivalent and are not shown. **B.** Rust’s increase due to the harvesting trajectories for each of the ripening scenarios (harvesting - control), for the five different planting densities (blue: asynchronous ripening, orange: synchronous ripening) and 30 repetitions. In all simulations, we started with 20% of infected plants at T_0_.

### 3.3. Medium size steps increase the proportion of infected networks of plants in the asynchronous scenario; the interaction between the proportion and size of infected networks explains the nonlinear behavior of rust increase

The proportion of infected networks of plants (where each pair of plants are 1.5 m apart or less) after harvest (# infected network AH/planting density) decreases as the planting density increases for both coffee ripening scenarios (subplot i in Fig. 5A, black circles in Fig. 5B). This makes sense since the connectivity between plants increases with density, making almost all the plants already infected before harvest due to the initial contact-mediated dispersal (gray circles in Fig. 5B). In the asynchronous ripening scenario, there are significantly more infected networks due to harvest trajectories than in the synchronous scenario (for planting densities higher than 500 plants/ha). This is due to the high proportion of medium sized steps between 1.5 and 8 m (58% of the steps) in trajectories generated under the asynchronous ripening (Fig.2), that potentially create new infected networks and contribute to the general infection (blue ribbon in Fig. 5C). In the synchronous scenario, 95% of steps in the trajectories are smaller or equal to 4 m (and 51% smaller than 1.5 m; Fig.2) and are more likely to disperse rust in the same contact-mediated networks, this is, in plants that are already infected (gray circles in Fig. 5B; orange ribbon in Fig.5C). Larger steps (>30 m) of trajectories in the asynchronous scenario, even if they connect distant zones of the plot, are not relevant for the infection as their proportion is very low (Fig. 5C).

**Fig. 5.**
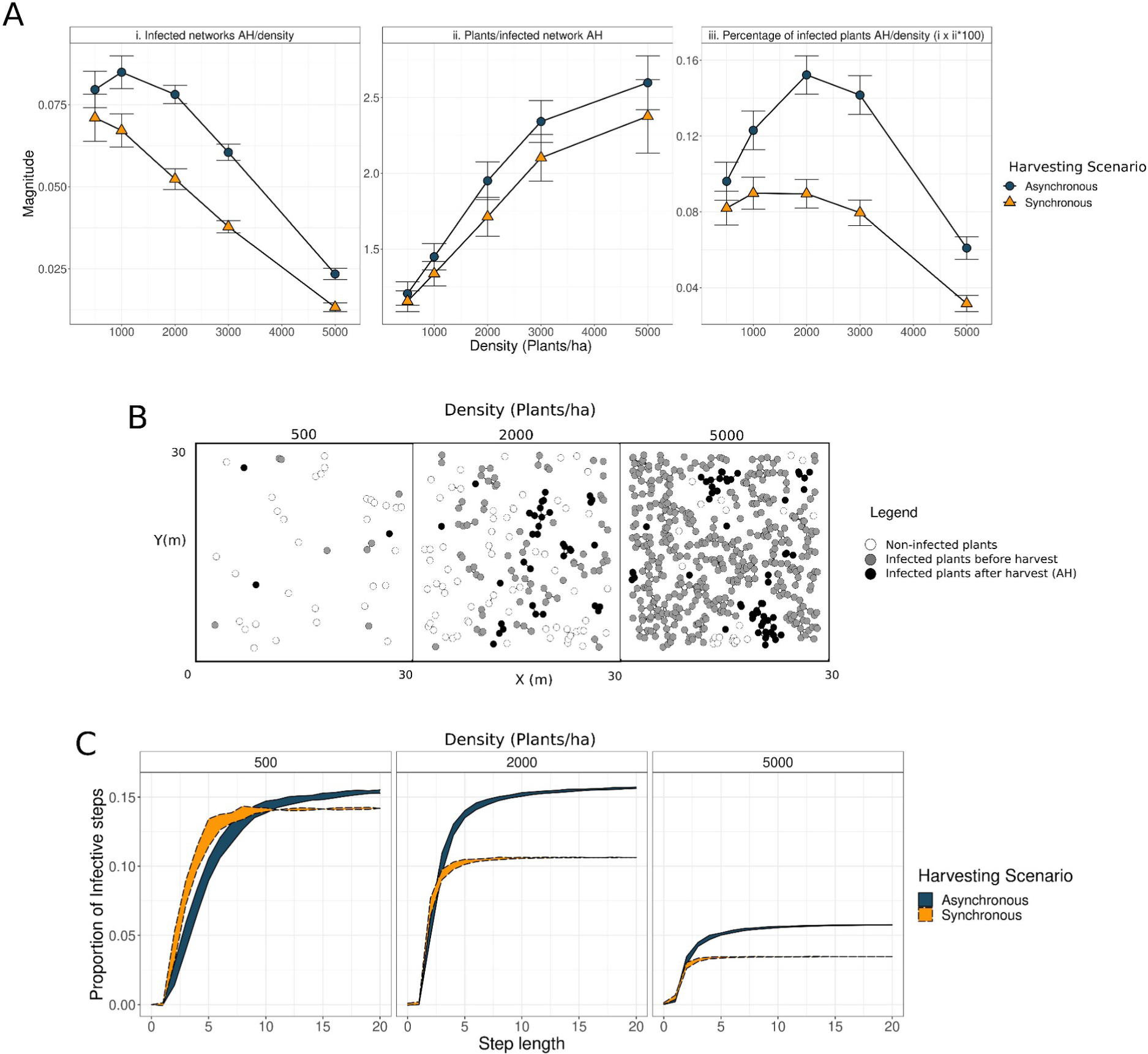
Proportion and mean size of infected networks after harvest and proportion of infective steps during harvest. **A.** We plot for each harvesting scenario (blue circle: asynchronous ripening, orange triangle: synchronous ripening) the proportion of infected networks after harvest (infected network AH/density; subplot i), their mean size (infected plants/infected network AH; subplot ii) and their product (100*infected plants AH/density; subplot iii). **B**. We show a 30 m2 zone of the plot in 500, 2000 and 5000 plants/ha densities to illustrate the non infected plants (white circles), the infected networks before harvest (gray circles) and the infected networks of plants after harvest and the cycles of contact-mediated dispersal (black circles). C. We show the proportion (mean ± standard deviation) of infective steps (steps that infected a new plant; yellow circles in Fig.1C) for each step length, three planting densities, and two harvesting scenarios.

On the other hand, the average size of new infected networks after harvest (# infected plants/infected network AH) increases with planting density as the connectivity increases (subplot ii in Fig. 5A, black circles in Fig. 5B). In this case, we found no significant differences between both harvesting scenarios. The estimated percentage of infected plants after harvest (% of plants AH/planting density; as the product of i and ii; subplot iii in Fig. 5A) follows a non-linear behavior in both scenarios equivalent to the observed increase of infection in Fig. 4. In this sense, the opposite behavior between the proportion of infected networks after harvest and their mean size might explain the nonlinear behavior in rust increase due to harvesting, with increasing planting densities. The higher rust increase in the asynchronous scenario is solely related to a higher proportion of infected networks after harvest due to the medium-sized steps.

For scenarios with very low initial infection (1%), almost no plants are infected before harvest, so the main driver to increase the general infection will be the size of the infected networks after harvest maximized at 5000 plants/ha (Fig. S1.4). On the contrary, at very high initial infections (50%), almost all the plants are infected before harvest, and only at very low densities (500 plants/ha), new plants can get infected by harvesting (Fig. S1.4).

## 4. DISCUSSION AND CONCLUSIONS

Many studies have stressed the relation between spatial structure of hosts, individual movement behavior, and pathogen transmission to predict and understand disease dynamics, though disease models with mechanistic representations remain rare (Manlove et al., 2022; White et al., 2018). Understanding the relationship between the spatial arrangement of the hosts and the different modes of dispersal of CLR is central to developing preventive strategies to reduce the impact of the pathogen on crop production (Hajian-Forooshani and Vandermeer, 2021).

Here we present a model to study the determinants of the movement of the workers in the field during harvesting and its relevance in CLR dispersal and infection at the plot level. Our results show that we can reproduce random walks, medium or large relocalizations by regulating the coffee ripening synchronicity and the scarcity of trees with berries at the end of harvest. These conditions could be *sufficient* to generate the different step lengths observed in the field. Relocalizations in deterministic walks on a random set of points have been documented for other systems (Brown et al., 2007) and simulated in rectangle domains (Boyer, 2008; Santos et al., 2007) or varying the size of the targets (Boyer et al., 2006). These studies present specific widths or target size uniformity ranges where the modeled workers fail to find closer unvisited points and “jump” to another zone of the area (Boyer, 2008; Boyer et al., 2006). Concerning the CLR impact, we found that increasing the planting density where only contact-mediated dispersal was present increases the final CLR at the plot level. This is in line with other studies (Hajian-Forooshani and Vandermeer, 2021; Zambolim, 2016; Mora Van Cauwelaert, et al. 2023) and emphasizes the importance of planting at lower densities to avoid big bursts of the pathogen (Ehrenbergerová et al., 2018). The harvest event magnifies the final CLR infection for all scenarios and densities. Its effect is more important for medium densities (2000 to 3000 plants/ha), which is a common density of coffee plots owned by small-scale farmers (Soto-Pinto et al., 2000). If densities are too high, harvesting becomes irrelevant as the plot is already infected. Other modes of dispersal, such as wind-mediated relocalizations from neighbor plantations, were not included (as in (Hajian-Forooshani and Vandermeer, 2021; Vandermeer and Rohani, 2014)). This assumption is reasonable for the rainy seasons where local contact is more relevant to build the initial epidemic (Boudrot et al., 2016). The different types of trajectories generated under the two coffee ripening scenarios changed the plot average CLR and varied nonlinearly with increasing planting density. Movements with medium and long steps in the asynchronous ripening scenario promote new foci of infection (Zadoks and Van Den Bosch, 1994) that eventually become newly infected networks with optimum size and number at planting densities between 2000 to 3000 plants/ha. In particular, the high proportion of medium-sized steps thrives the infection due to its high probability to infect distant coffee networks. This movement is common in plots with asynchronous ripening where workers jump from one row to the other to maximize harvest (Mora Van Cauwelaert et al., 2023b). In this sense, the spatial heterogeneity in coffee maturation creates fragmented host spatial patterns that interact with the movement of harvesters and lead to CLR outbreaks at landscape levels (as suggested by White et al., 2018). It is important to note that we modeled a random arrangement of the plants in the lattice. This assumption is likely valid for old plantations and shaded coffee managements (Moguel and Toledo, 1999) but should be revisited for aggregated or uniform patterns.

While we observe an increase in CLR in the asynchronous scenario, our results do not imply that asynchronous ripening leads *per se* to higher rust. On the one hand, the long steps are not generated due to the asynchronous ripening alone, but mainly due to the scarcity of closer trees at the end of harvest. In fact, when we run the synchronous scenario in a scarcity scenario, longer steps also appear (data not shown). This would also be the case for large-scale landlord owned plantations where the workers are paid by daily harvest (Jiménez-Soto, 2021); even if the following plant is hundreds of meters distant, it becomes necessary for them to reach it within the same day (Mora Van Cauwelaert et al., 2023b). In these plantations it is also common to force the workers to harvest at the end of the season when only few scattered trees still have berries (Mora Van Cauwelaert et al., 2023b). This would promote medium-sized steps, the main drivers of higher plot level infection. On the other hand, the simulated trajectories in the two coffee ripening scenarios suppose otherwise similar conditions on the plot. But in the field, asynchronous ripening is associated with characteristics that change these conditions and that can reduce the general rust impact. For example, the susceptibility of coffee plants to rust increases with fruit charge (Salgado et al., 2008). Hence, the asynchrony in the maturation of berries due to erratic rainfalls, or simply due to different coffee varieties (DaMatta et al., 2007), generates a mixed spatial pattern of resistant and susceptible plants that could reduce rust dispersal. Besides, shade trees in the plot form different patches of maturation (and the potential for long steps during harvest) but also create barriers to wind-mediated rust dispersal (Gagliardi et al., 2020).

Our study aims to spur discussion on practices that can reduce the impact of harvesting and specific trajectories, in scenarios where one can benefit from asynchronous maturation of berries and shaded plantations (Ehrenbergerová et al., 2018; Liebig et al., 2019; Soto-Pinto et al., 2000). This is crucial as almost all the plantations have or will have this kind of ripening (DaMatta et al., 2007; Masarirambi et al., 2009), especially in the predicted climate scenarios of increasingly erratic rainfalls (Avelino et al., 2015; Huang et al., 2012; Rana et al., 2014). Along with planting different genetic varieties (Silva et al., 2006) or having other species of trees inside the plot and between them (Avelino et al., 2022), our results show the importance of reducing medium to long-distance movements during the same day by avoiding harvesting at the end of the harvest season when trees ripening asynchronicity and CLR levels are higher (Avelino et al., 1991; Mora Van Cauwelaert, 2023b); or simply skipping rust-infected trees and working fewer hours a day, even if this represents a short term decrease in harvested berries. These practices presuppose the right to decide how and when to harvest the coffee plantations.

We hope this mechanistic model will contribute to educating our intuition on the relationships between host spatial disposition and dispersal, and it will help us build working hypotheses for an agroecological approach to pest management that does not rely solely on the control of the infection stage of the pathogen, but also on preventive measures that focus on the dispersal of the fungus.

## ACKNOWLEDGMENTS

EMVC is a doctoral student from the Programa de Doctorado en Ciencias Biomédicas, Universidad Nacional Autónoma de México, and has received CONACyT scholarship 686776 and support from the Programa de Apoyo a los Estudios del Posgrado (PAEP) for his research stay at the University of Michigan (USA). MB acknowledges financial support from UNAM-DGAPA-PAPIIT (IN207819). JV acknowledges financial support from the US National Science Foundation, grant number DEB-1853261. EMVC thanks the people from La Parcela and Perfectomeer lab for their great ideas and friendship. EMVC and MBK are also thankful to Gustavo Bautista, Gabriel Domínguez, Elisa Lotero and Cecilia González for the discussions and help during the fieldwork. The authors thank Denis Boyer and Eugenio Azpeitia for their valuable feedback on previous versions of the manuscript and Rodrigo García Herrera for his technical support.

## 6. CRediT

**EMVC**: Conceptualization, Methodology, Software, Validation, Formal Analysis, Investigation, Writing-Original Draft, Visualization. **KL**: Conceptualization, Methodology, Writing - Review & Editing. **ZHF**: Conceptualization, Methodology, Software, Writing - Review & Editing. **JV:** Conceptualization, Methodology, Writing - Review & Editing, Supervision, Funding acquisition. **MB:** Conceptualization, Methodology, Writing - Review & Editing, Supervision, Project administration, Funding acquisition

## Notes

### Competing Interest Statement

The authors have declared no competing interest.

### Summary of Updates

We updated all the figures. This new version has a clearer and more precise interpretation of the results. We also added some changes in the methods and model. Our new results point to middle-sized steps as the main factor to rust increase at plot level. We also added new supplementary figures and tables.

