## supplementary figures and tables for "Interplay between harvesting, planting density and ripening time affects coffee leaf rust dispersal and infection"

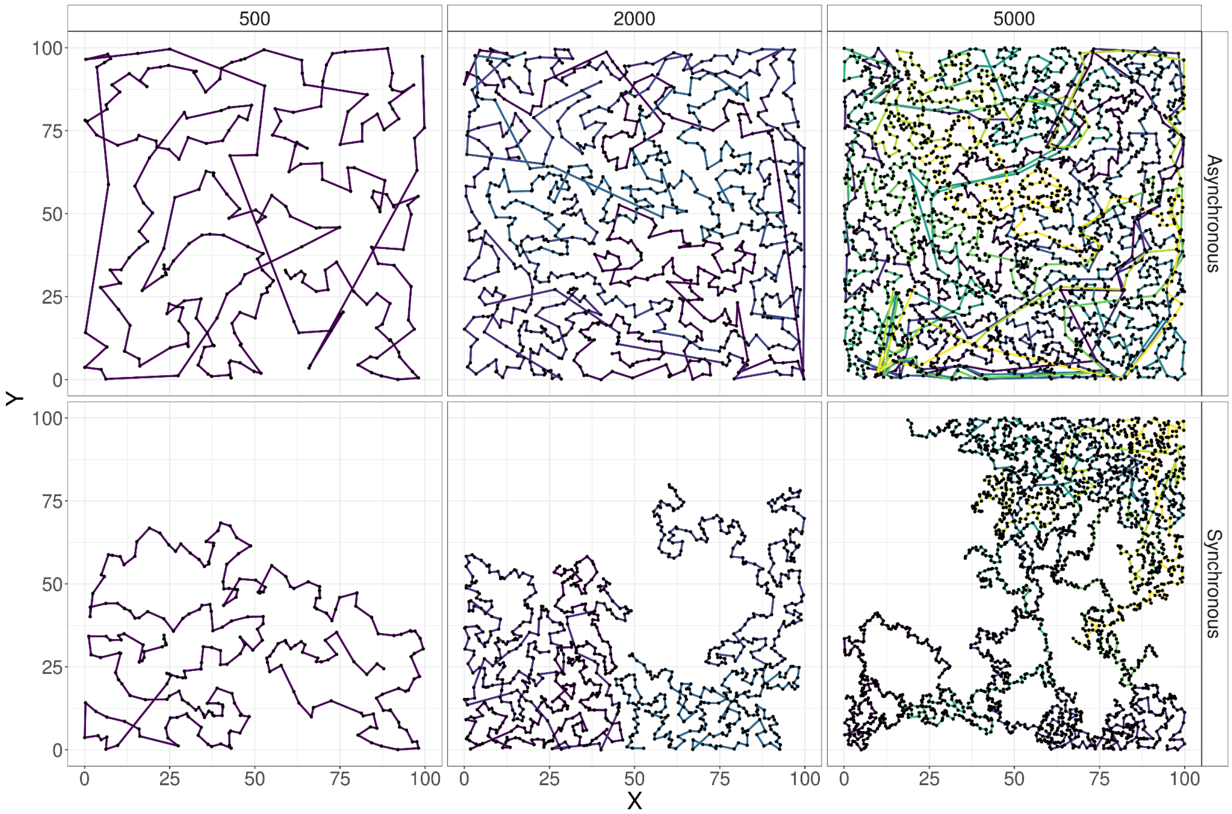


**Figure S1. 1. Simulated trajectories under two coffee ripening scenarios and three planting densities.** The columns represent the three planting densities (plants/ha) and each row represents one ripening scenario. Colors correspond to each of the workers (1, 2 or 5 workers for 500, 2000 and 5000 plants). In all scenarios, the simulations were run until half of the plants were harvested (N/2). The trajectories of N= 1000 and N= 3000 were qualitatively the same and are not shown.

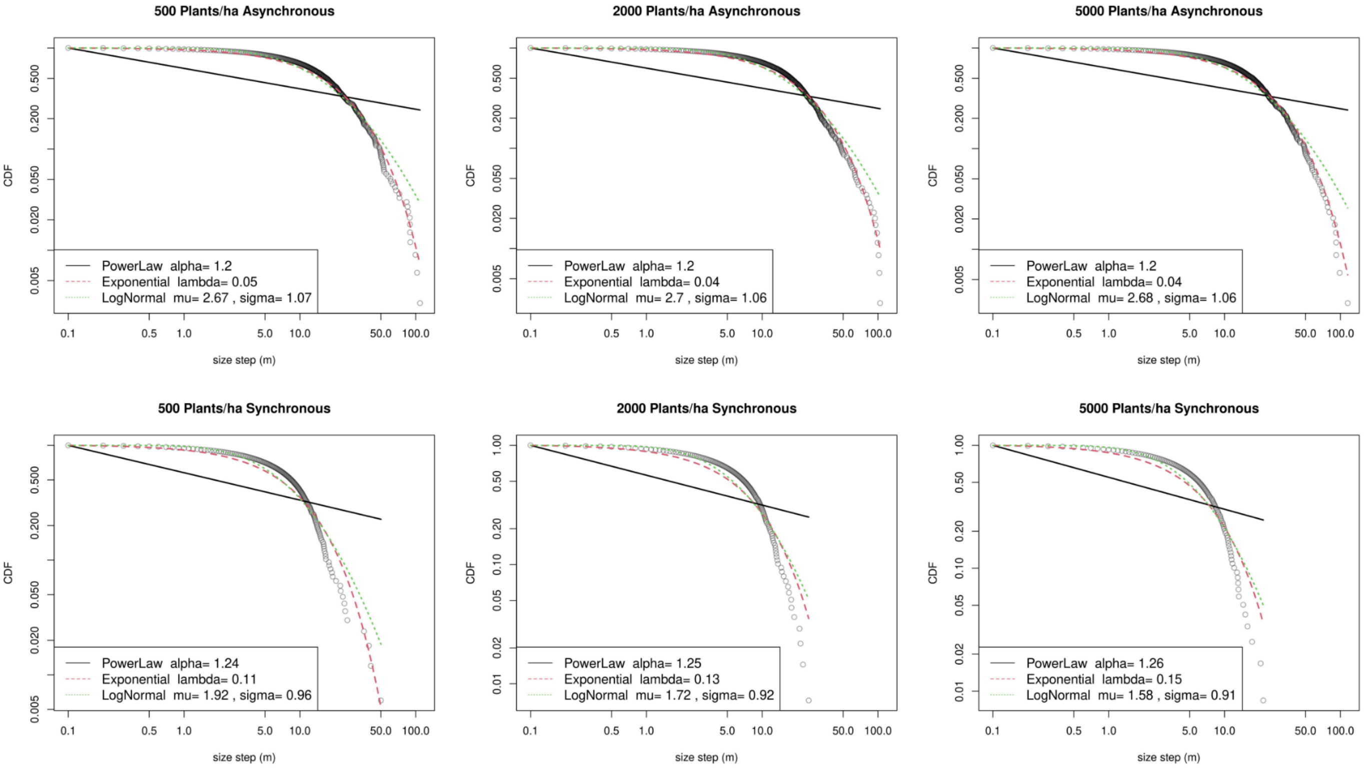


**Figure S1. 2. Cumulative distribution function and fitting of three models for the distribution of steps under two coffee ripening scenarios and three planting densities.** The data for each scenario is presented with empty circles, and the three models are presented as lines (power law: black continuous, exponential: red dotted, lognormal: green dotted line). The characteristic parameters are presented in the small box of each plot.

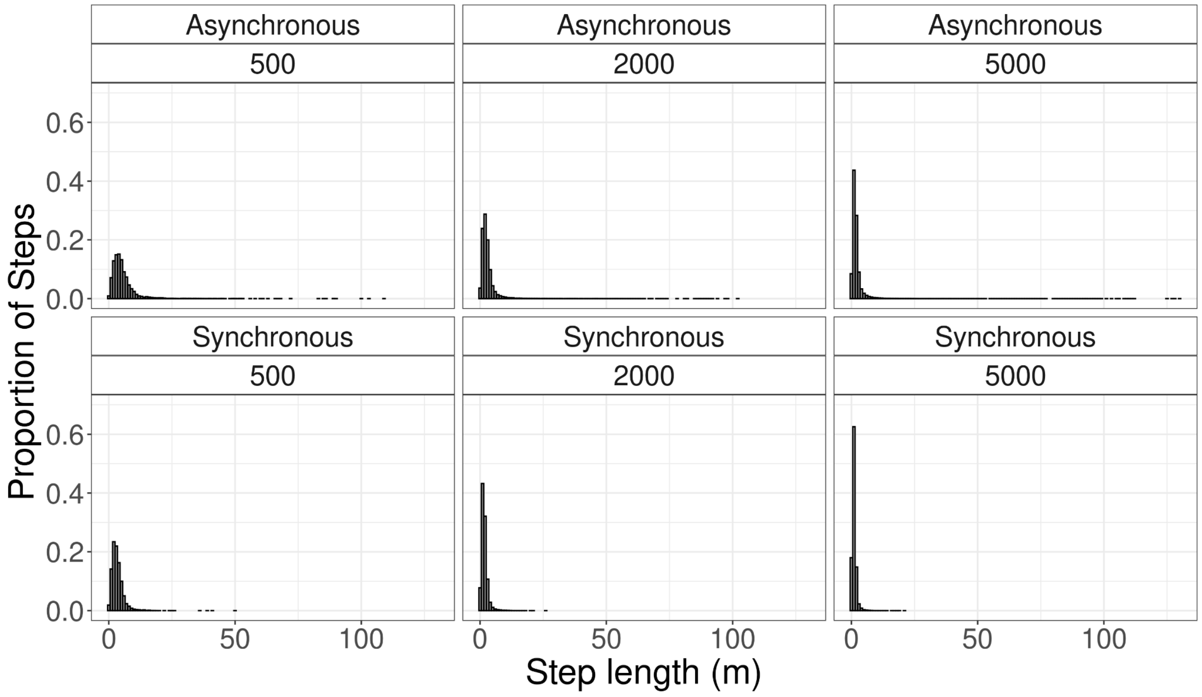

**Figure S1. 3. Step length distribution for three densities and two coffee ripening scenarios.**

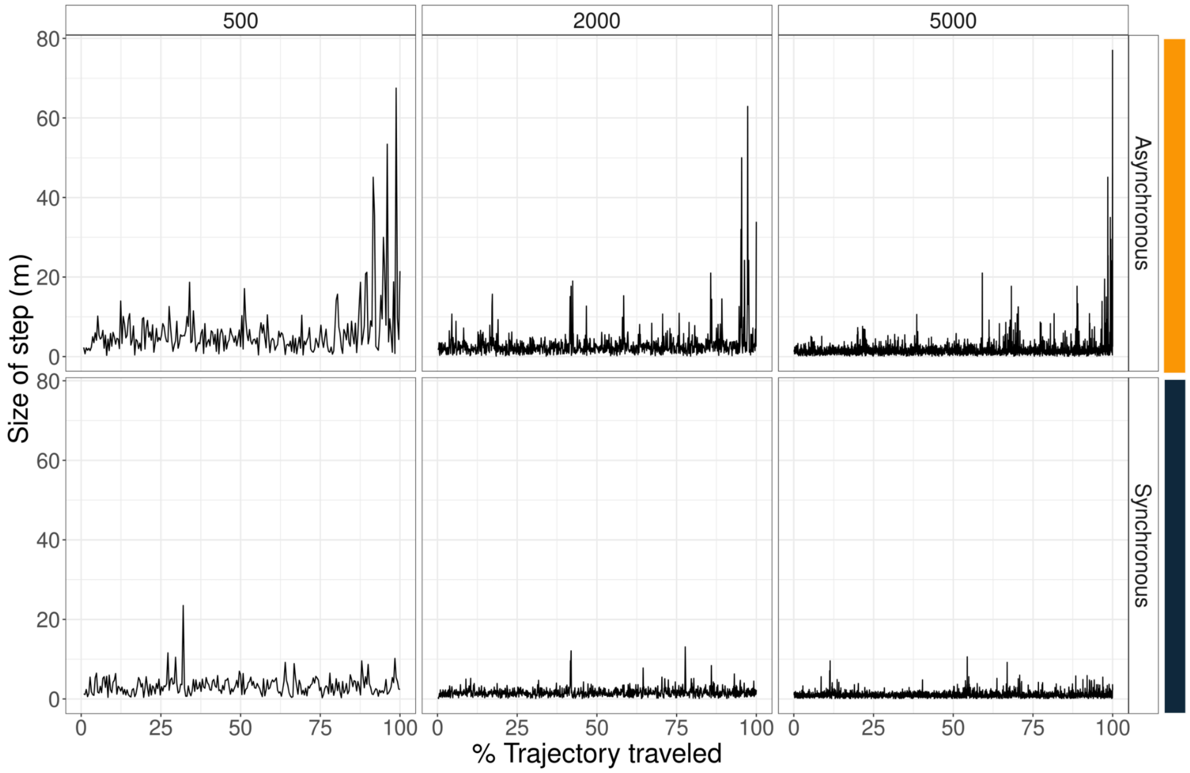


**Figure S1. 4. Size of step (in m) in relation to the percentage of the trajectory traveled ((number of trees visited/total trees visited) x100), under two coffee ripening scenarios and three planting densities.** The % of trajectory traveled represents the number of current visited trees normalized by the maximum number of visited trees per scenario. The columns represent the three planting densities (plants/ha) and each row the coffee ripening scenario. In all scenarios, the simulations were run until half of the plants were harvested (N/2). The dynamics of N= 1000 and N= 3000 were qualitatively the same and are not shown.

**Table S1.1.** Central tendency values of the distribution of steps for five different plating densities and two coffee ripening scenarios. The mean distance, mode and median are in meters.

| Density (plants/ha) | Harvesting Scenario | Median | Mean | SD | Mode |
|---|---|---|---|---|---|
| 500 | Asynchronous | 4 | 5.77 | 6.47 | 4 |
| | Synchronous | 3 | 3.36 | 2.40 | 2 |
| 1000 | Asynchronous | 3 | 4.26 | 5.79 | 2 |
| | Synchronous | 2 | 2.37 | 1.67 | 2 |
| 2000 | Asynchronous | 2 | 3.23 | 5.17 | 2 |
| | Synchronous | 1 | 1.69 | 1.25 | 1 |
| 3000 | Asynchronous | 2 | 2.72 | 4.71 | 1 |
| | Synchronous | 1 | 1.40 | 1.12 | 1 |
| 5000 | Asynchronous | 1 | 2.21 | 4.58 | 1 |
| | Synchronous | 1 | 1.08 | 0.86 | 1 |

**Table S1.2.** Cumulative proportion of steps per step length. For each scenario (five planting densities and two harvesting modes), we show the cumulative proportion of steps lower than 1.49 m (rounded to 1 m), and the step length when the cumulative proportion reached 0.95.

| Number of plants | Harvesting Scenario | Cumulative proportion | Step length |
|---|---|---|---|
| 500 | Asynchronous | 0.08 | 1 |
| | | 0.95 | 14 |
| | Synchronous | 0.16 | 1 |
| | | 0.95 | 7 |
| 1000 | Asynchronous | 0.16 | 1 |
| | | 0.95 | 10 |
| | Synchronous | 0.30 | 1 |
| | | 0.96 | 5 |
| 2000 | Asynchronous | 0.27 | 1 |
| | | 0.95 | 8 |
| | Synchronous | 0.51 | 1 |
| | | 0.97 | 4 |
| 3000 | Asynchronous | 0.37 | 1 |
| | | 0.95 | 7 |
| | Synchronous | 0.64 | 1 |
| | | 0.96 | 3 |
| 5000 | Asynchronous | 0.52 | 1 |
| | | 0.95 | 5 |
| | Synchronous | 0.81 | 1 |
| | | 0.95 | 2 |

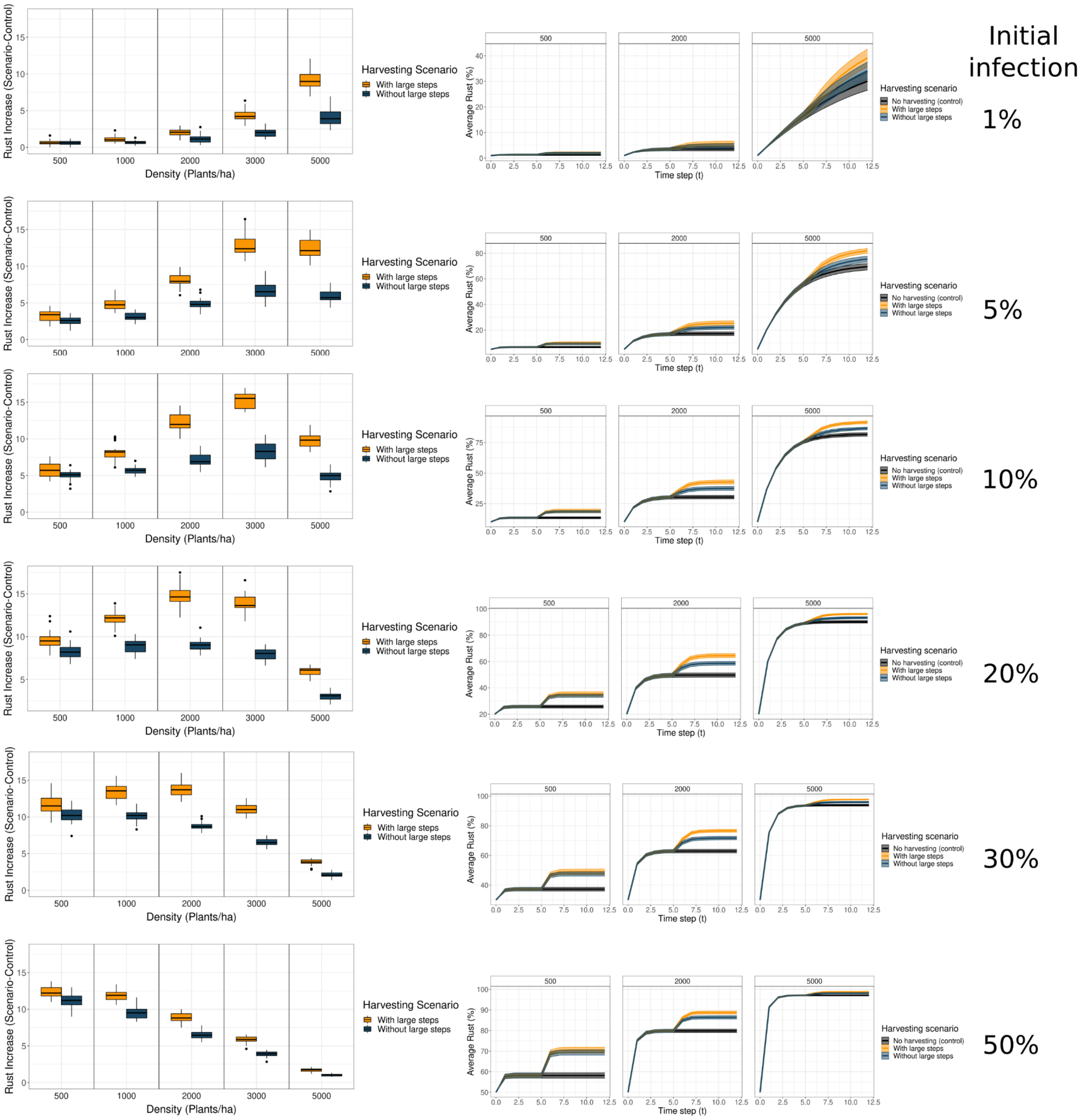


**Fig. S1.5.** Effect of different initial infection conditions (1 to 50%) on the rust's increase due to the harvest (left column) for each of the trajectories (harvesting scenario - control) for the five different planting densities (orange box: with large steps, blue: without large steps) and on the full time series of the average rust for three planting densities. The percentage in the right indicates the initial infection.
